# Sponges of the Lower Greensand Group (Lower Cretaceous) of England; a revision

**DOI:** 10.1101/2024.06.28.601142

**Authors:** Consuelo Sendino, Stephen Kershaw

**Affiliations:** Collections Department, National Museum of Natural Sciences, Jose Gutierrez Abascal 2, 28006, Madrid, Spain; Science Group, Natural History Museum, Cromwell Road, London, SW7 5BD, UK; Department of Life Sciences, Brunel University London, Uxbridge, UB8 3PH, UK

**Keywords:** Lower Greensand Group, Faringdon Sponge Gravel Member, Porifera, taxonomy, calcareous sponges

## Abstract

Sponges of the Lower Greensand Group (LGS) are well preserved, and occur in sediments of a sandy matrix. Abundant in the Faringdon Sponge Gravel Member (FSG), these sponges, mostly Calcareans are found in Oxfordshire, with notable preservation at Little Coxwell quarries. Classical researchers described sponges and spicules from the LGS, including Lhuyd (considered to have been the first to publish illustrations of LGS sponges), Sharpe, Sowerby and Parkinson. In addition to the FSG, the Folkestone, Hythe, and Atherfield Clay formations within the LGS also contain sponge remains, including spicules as well as whole sponge fossils. These sponges include mostly samples from traditional sponge class Calcarea and a taxon of Hexactinellida; Altogether, the sponge assemblage developed in warm seas of the Lower Cretaceous, and display diverse shapes of sponge bodies and robust spicules. This study provides descriptions of common species following updated Porifera classification and recent sponge taxonomy research, illustrated with specimens from the Natural History Museum, London (NHM) and British Geological Survey (BGS) collections. The following taxa are recorded and described: 1) Calcareans: *Barroisia anastomosans* (Parkinson, 1811), *Barroisia clavata* (Keeping, 1883), *Barroisia irregularis* (Hinde, 1884), *Dehukia crassa* (de Fromentel, 1861), *[Elasmoierea] faringdonensis* (Mantell, 1854), *[Elasmoierea] mantelli* (Hinde, 1884), *Peronidella gillieroni* (Loriol, 1869), *Peronidella prolifera* (Hinde, 1884), *Peronidella ramose* (Roemer, 1839), *Oculospongia dilatate* (Roemer, 1864), *Tremospongia pulvinaria* (Goldfuss, 1826), *Raphidonema contortum* (Hinde, 1884), *Raphidonema porcatum* (Sharpe, 1854), *Raphidonema farringdonensis* (Sharpe, 1854), *Raphidonema macropora* (Sharpe, 1854), *Raphidonema pustulatum* Hinde, 1884, *Endostoma foraminosa* (Goldfuss, 1829); 2) Hexactinellids: *Lonsda contortuplicata* Lonsdale, 1849.

## 1. Introduction

Sponges with other organic remains, mostly marine, from the Lower Greensand Group (LGS) have been described from the beginning of 19th century. In most cases they are well-preserved, presumably as a result of their hypercalcified nature (the sponges secreted a basal skeleton of calcium carbonate, as a secondary skeleton as they grew. The sponges are found mainly in poorly consolidated sandy sediment. Amorphozoae, as sponges were named at the beginning of their study, can be found abundantly in the Faringdon Sponge Gravel Member (FSG, see Hopson *et al*. 2008, table 5), within the Faringdon Sand Formation. The FSG is thus stratigraphically positioned in the upper part of Lower Cretaceous strata. Sponges stand out in Oxfordshire (former historic Berkshire), at the Faringdon outcrop, where they can be found in coarse pebbly and cross-stratified sands 10 m thick (Krantz 1972; Hesselbo *et al*. 1990) and also at Little Coxwell quarries, with sponge assemblages with spectacular preservation. This member comprises predominately calcareous sponges of littoral facies. Sharpe (1854) was one of the first palaeontologists to describe sponges from this member, and Hinde (1884) made a taxonomic study of the sponges of this group deposited at the British Museum (Natural History) [former Natural History Museum]. But, undoubtedly, the first one to pay attention to sponges of the FSG was Lhuyd (1699) who cited the Coxwell Sponge-Gravel pits and the Faringdon outcrop. He published the first recorded illustrations of sponges from the LGS, including ‘*Endostoma foraminosa’* (Lhuyd 1699: pl. 18: fig. 1522). Other classical researchers were Sowerby (1811) who mentioned sponges and sponge spicules from the FSG, and Parkinson (1822) and Mantell (1839, 1844, 1848, 1854) who described new species from this stratum.

Other formations in this group with sponges as well are the Folkestone (FF), Hythe (HF) and Atherfield Clay (ACF) Formations. The FF sponges are normally represented by spicules (Casey 1961), but at Coxbridge Pit they have a standard high state of preservation. Spicules have been found in siliceous rocks of the FF in an excavation at the at Baker’s Gap (East Cliff), and Cop Point in the third and fourth divisions respectively of the FF by Price (1874). Casey (1961) also reported sponge spicules at Sandling Junction Sandpit and the Iguanodon Quarry in the same formation as well. HF contains numerous sponge spicules at the Godstone area of Surrey, in the chert beds of Tilburstow and Haslemere in which Hinde (1885) found spicules belonging to 28 taxa. Finally, spicules and few sponges have been reported from ACF in Atherfield and Sandown, on the Isle of Wight.

The LGS sponges, mostly FSG, lived in the warm shallow Lower Cretaceous waters in England that was then at a latitude of around 40 degrees north (e.g. Scotese, 2015, see Fig. 1) and are characterised by mostly irregular shapes, typical of shallow waters, containing robust spicules and having a short base to attach to the seabed, or they encrusted on rocks or other sponges. Most of these are sponges of the class Calcarea, but there are few hexactinellids and demosponges of deeper waters that have not been studied here. This article includes the description of the most common British species (17 calcareous sponges and one hexactinellid) of the Lower Greensand and may be regarded as characteristic. For this we have followed the most updated Porifera classification (Finks *et al*. 2004a), the most recent scientific articles on sponge taxonomy (De Laubenfels, 1955; Delamette et al., 1986; Finks, 2004; Hurcewicz, 1975; Masse and Termier, 1992; Senowbari-Daryan et al., 2011; Vacelet, 1979) and illustrated the species with specimens housed at the Natural History Museum, London, (NHM) and the British Geological Survey (BGS).

**Figure 1.**
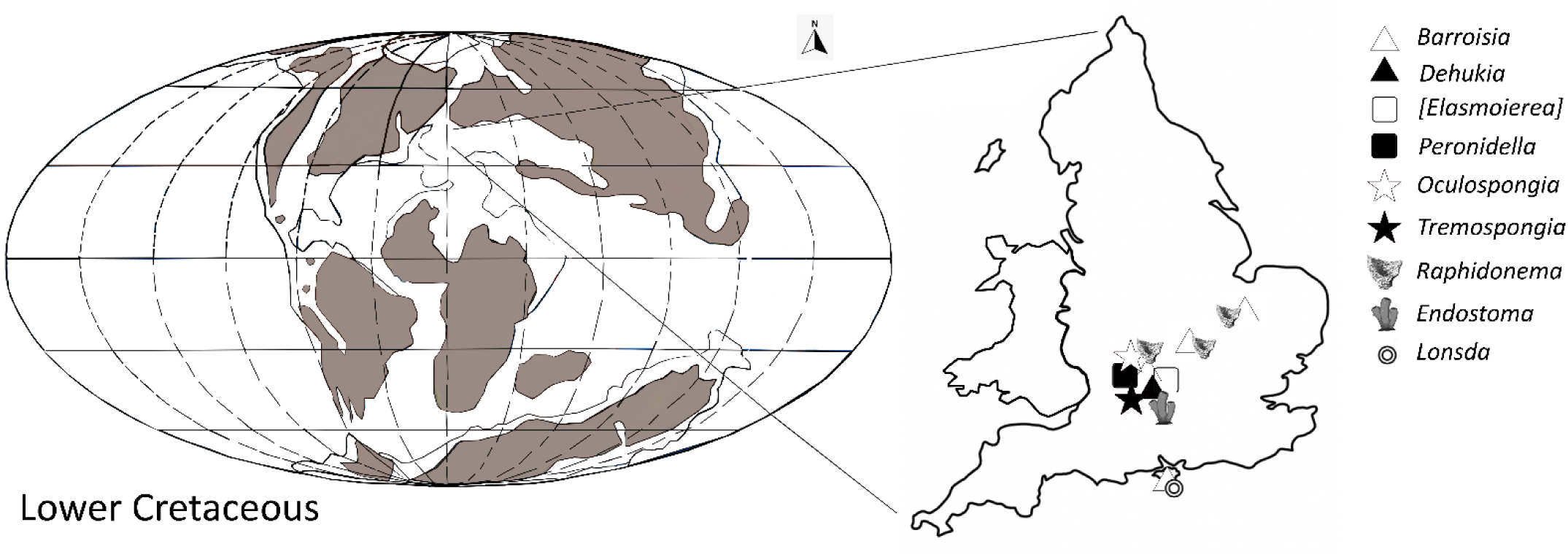
Continent and oceanic distribution during LGS, Lower Cretaceous, along with genera occurrences of the British LGS porifera forms continental reconstruction redrawn from Scotese, 2015).

## 2. Material and methods

We have studied British specimens housed in official institutions such as the Natural History Museum, London (NHM) and the British Geological Survey (BGS).

In order to understand the sponge descriptions, below are defined the most used morphological terms in this study, using the sphinctozoan-grade sponge in Fig. 2 to emphasise some terms.

**Figure 2.**
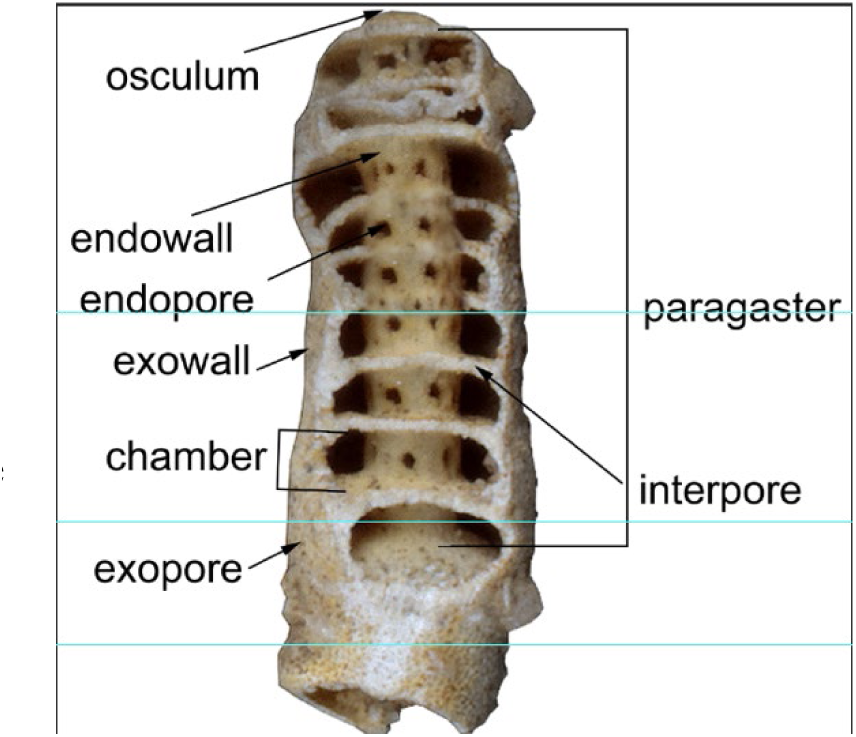
Sphinctozoan-grade sponge with main morphological elements

*Apopore:* Exhalant pore.

*Clavate*: form gradually thickening toward the distal end, club-shaped.

*Cloaca*: large, central exhalant cavity, without digestive function. Also termed e.g. spongocoel or paragaster.

*Dermal layer* (*skeletal*): Differentiated peripherical top layer, specialized or not.

*Diactine:* Spicule with two rays.

*Ectosome*: Cortical part, directly beneath the exopinacodenn or outer layer of a sponge, characterized by the absence of choanocyte chambers and extending across the outer ends of inhalant canals.

*Endopore*: Pore that pierces the inner wall of a chamber (Fig. 2).

*Endowall*: Wall of a central exhalant tube or cloaca (Fig. 2).

*Equiradiate*: Equal rays.

*Exowall*: External skeleton of a chamber (Fig. 2).

*Exopores*: Pore that pierces the outer wall of a chamber (Fig. 2).

*Filiform*: Thread or filament-like shape.

*Interpore*: Pore that pierces the wall between chambers (Fig. 2).

*Nodose excrescence*: Knotty distinct outgrowth.

*Osculum*: Large aperture of the exhalant canal or paragaster.

*Paragaster*: Large internal, central exhalant cavity, without digestive function (Fig. 2).

*Paratangential dermal spicule*: Dermal spicule near to being tangent.

*Parenchymal*: regarding endosome or inner part, in middle layer of body wall, between dermal and atrial layers; where the spicules are located.

*Tetraradiate*: Spicule with four rays

*Trabecular (network)*: Microstructure pillar-like filling skeleton.

*Triradiates*: Spicule with three rays.

*Tylostyle*: Monaxon, or spicule in which ray grow along a single growth axis, knobbed at one end, sharply pointed at the other.

*Tylote*: Monaxon, or spicule in which ray grow along a single growth axis, knobbed at both ends.

## 3. Systematic palaeontology

### SPECIES DESCRIPTIONS

> Phylum PORIFERA Grant, 1836
>
> Class CALCAREA Bowerbank, 1864
>
> [=class Calcispongea De Blainville, 1834; *ex* order Calcispongiae De Blainville, 1834; Calcarosa Haeckel, 1872; Megamastictora Sollas, 1887]
>
> Subclass CALCARONEA Bidder, 1898
>
> Order SPHAEROCOELIIDA Vacelet, 1979
>
> Family SPHAEROCOELIIDAE Steinmann, 1882
>
> Genus BARROISIA Munier-Chalmas, 1882

#### Diagnosis

Erect, conicocylindrical branching tubes formed of irregular chambers which may be segmented only internally. Conical paragaster occupies one-third of the sponge diameter. Exowall netlike with subpolygonal, substellate exopores. Partitions between chambers gently arched, mostly upwards distally, chambers low, with polygonal interpores. Endowall continuous, with horizontal whorl of large, circular endopores in each chamber. Exowall made of equiradiate triradiates on the inner layer parallel to wall and penicillately arranged tylostyles, tylotes on the outer layer, both embedded in finely fibrous groundmass.

#### Remarks

Provisionally included in the Sphaerocoeliidae due to the existence of triradiates.

#### Type species

*Tubipora anastomosans* Parkinson, 1822.

> *Barroisia anastomosans* (Parkinson, 1822)
>
> Figures 3A-D
>
> 1822 *Tubipora anastomosans* Parkinson, 70-71, pl. 9: fig. 10
>
> 1839 *Tubipora anastomosans* sensu Mantell, 560: fig. 105.3
>
> 1843 *Verticillipora anastomosans* (Mantell) in Morris, 46
>
> 1844 *Verticillipora anastomosans* (Mantell), 289-290, fig. 55.4
>
> 1848 *Verticillipora anastomosans* (Mantell), p. 636, fig. 139.3
>
> 1848 *Verticillipora anastomosans* (Mantell) in Bronn, 1364.
>
> 1854 *Verticillipora anastomosans* (Mantell), 273, figs 70.4, 72.3.
>
> 1854 *Verticillipora anastomosans* (Mantell) in Sharpe, 195, pl 5: figs 1a-1e.
>
> 1874 *Verticillites anastomosans* (Mantell) in Davey, 13-14. pl. 7: fig. middle; pl. 8: fig upper.
>
> 1882 *Barroisia anastomosans* (Mantell) in Munier-Chalmas, 425.
>
> 1882 *Barroisia anastomans* (Mantell) in Steinmann, 164, pl. 8: figs 1-1a.
>
> 1883 *Verticellites anastomosans* (Mantell) in Keeping, 145.
>
> 1883 *Verticillites anastomans* (Mantell) in Carter, 27.
>
> pars 1884 *Tremacystia anastomans* (Mantell) in Hinde, 175, pl. 34: fig. 4a-c.
>
> 1890 *Barroisia anastomosans* (Mantell) in Steinmann and Döderlein, 72-73, fig. 68A-C.
>
> 1905 *Tremacystia anastomans* (Mantell) sensu Hinde in Davey, 17-18, pl. 1: fig. 2; pl. 2: fig. 1.
>
> 1914 *Barroisia anastomans* (Mantell) in Rauff, 84-103, pl. 1: figs 1-5; pl. 2: 6-11.
>
> 1914 *Barroisia anastomosans* (Mantell) in Douville, 399, pl. 12: figs 4-5.
>
> pars 1986 *Barroisia anastomosans* (Mantell) in Delamette and Termier, 314, pl. 2: figs 1-6, 9-12, text fig. 4.
>
> 1992 *Barroisia anastomosans* (Mantell) in Masse and Termier, 96, pl. 6: figs 1-4.

#### Description

Bush-like colony with cylindrical tubes composed of superposed chambers. The cylindrical forms are ca. 30-60 mm high with approximate diameter of 5-7 mm. The width of the conical paragaster is less than one-third of this diameter. These merge at their base, but are usually free distally (Fig. 3B). Lack of paragaster at the proximal area (Fig. 3D). Outer surface may be slightly constricted at the chamber partition level, but without external segmentation (Fig. 3A). Chamber height with an average of 2.3 mm. Endowall continuous with horizontal whorl of large, coarse, circular endopores in each chamber, 0.4 mm thick. This whorl extends downwards and upwards making the wall firm. Exowall with subpolygonal exopores and triradiate on the inner layer, thickness approximately 0.4 mm. With radial canals perpendicular to exowall (Fig. 3C). Subcircular apopores arranged at the whorl. Chambers with weak crenulations on their floor and roof (Fig. 3C). Internal segmentations with almost 1 mm thick.

**Fig. 3A-D.**
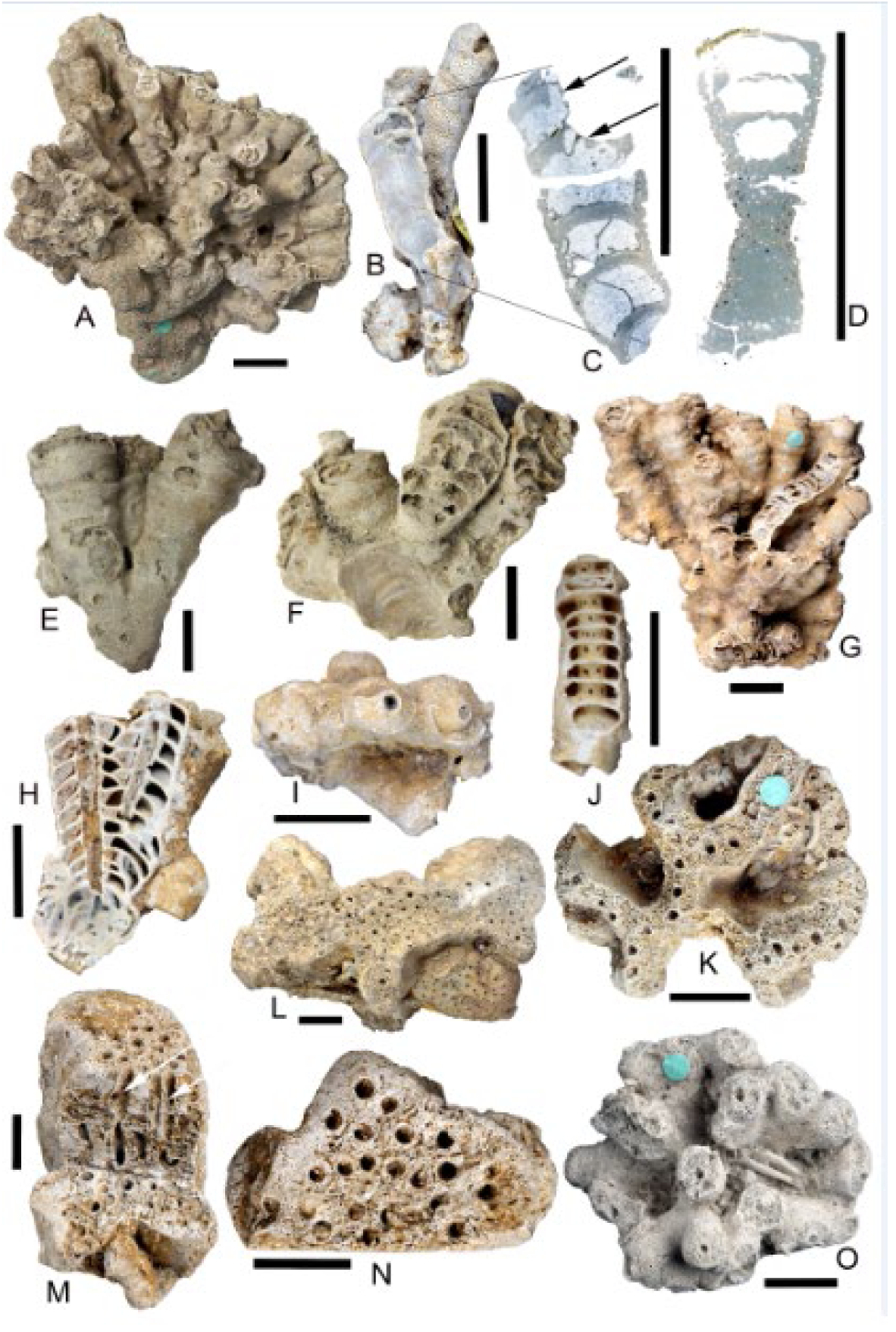
*Barroisia anastomosans* (Parkinson, 1822) (A. NHMUK PI P 3280 from Faringdon, Oxfordshire. B-D. NHMUK PI P 3264 from Upware, Cambridgeshire). A. Typical bush colony. B. Longitudinal section through chambers. C-D. Cross sections of a cylindrical form seen on B with visible internal partitions. C. Radial canals seen with vesicular internal filling, beginning of the paragaster partially seen at the last two partitions on the figure (black arrows). D. Base and first chambers of an individual. Wall detail with exopores and endospores. E-F. *Barroisia clavata* (Keeping, 1883) (E. Holotype BGS SM B 26236. F. Paratype BGS SM B 26237 from Upware, Cambridgeshire). E. Lateral view of colony with three forms in contact. F. Longitudinal section through chambers. 1G-1J. *Barroisia irregularis* (Hinde, 1884) (G. Holotype NHMUK PI P 3271. Paratypes: H-I. NHMUK PI P 2188 and J. NHMUK PI S 4619. All from Faringdon, in Oxfordshire). G. General view of the holotype with typical branching form. H. Longitudinal section of two attached tubes with interconnected chambers. I. View from the upper part, with protruding oscula. J. Endowall seen. K. *Dehukia crassa* (de Fromentel, 1861), NHMUK PI P 3273 from Faringdon, in Oxfordshire, seen from the upper part with the typical anastoming wall and oscula arrangement. L. *[Elasmoierea] faringdonensis* (Mantell, 1854), NHMUK PI S 4459, from Faringdon, Oxfordshire, fan-shape on the upper part, oscula with irregular arrangement, and attached to *Raphidonema pustulatum* Hinde, 1884. M-N. *[Elasmoierea] mantelli* (Hinde, 1884), paratype NHMUK PI P 3491, from Faringdon, in Oxfordshire. M. Longitudinal section with cylindrical exhalant canals exposed. N. View from the upper part with oscula seen. O. *Peronidella gillieroni* (Loriol, 1869), NHMUK PI OR 10202, from Faringdon, Oxfordshire, general view. Scale bar 10 mm.

**Fig. 4A-C.**
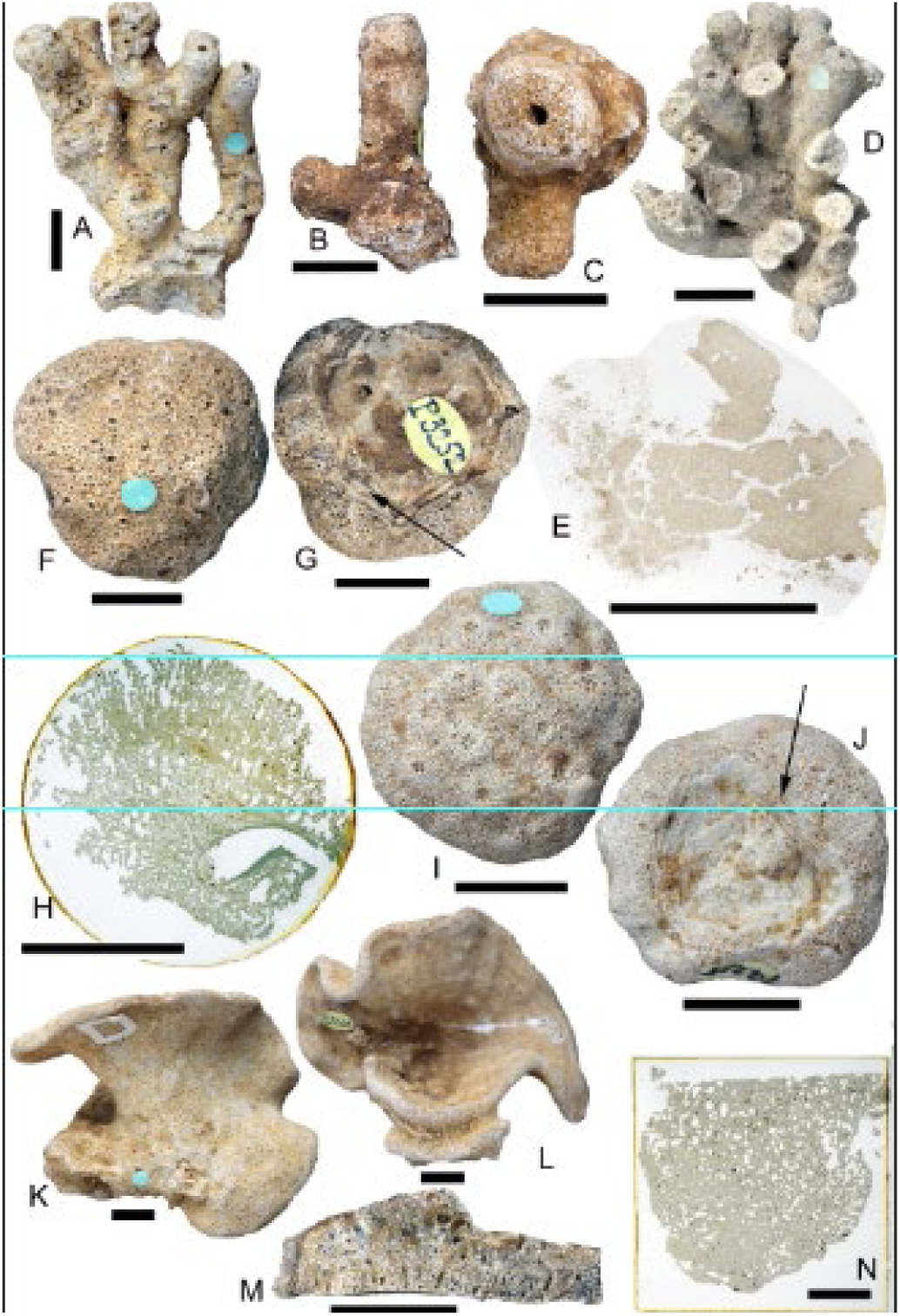
*Peronidella prolifera* (Hinde, 1884) from Faringdon, Oxfordshire. A. Holotype, NHMUK PI P 3277, colony general view. B-C. NHM UK PI P 4188. B. Forms bifurcating. C. Form summit with central circular osculum. D-E. *Peronidella ramosa* (Roemer, 1839), from Faringdon, Oxfordshire. D. NHMUK PI S 4542, general view. E. NHMUK PI P 2174, skeletal fibres in tangential section. F-H. *Oculospongia dilatata* Roemer, 1864 from Faringdon, Oxfordshire. F. General view from the upper part, oscula irregularly distributed. G. General view from the base, with concentric imperforate dermal layer (with an arrow). H. Vertical section with radiating fibres connected transversally. I-J. *Tremospongia pulvinaria* (Goldfuss, 1826), NHMUK PI OR 10215 from Faringdon, Oxfordshire. I. Upper view showing osculum clusters with asterisk shape, each one formed of 5 or 6 openings at a slightly higher level than the rest of the surface. J. Base showing the concentric imperforate dermal layer (with an arrow). K-N. *Raphidonema contortum* Hinde, 1884 from Faringdon, Oxfordshire. K. Paratype NHMUK PI P 3246, general view of one side of the cup shape with oscula seen. L. Same specimen as K seen from the other side with view of the cup shape. M. Paratype, NHMUK PI P 2178, wall cross section. N. Paratype, NHMUK PI S 8575, microstructure seen in thin section. Scale bar 10 mm.

#### Occurrence

FSG, LGS; Aptian, Lower Cretaceous of England, from Faringdon, in Oxfordshire; Upware, in Cambridgeshire; Little Brickhill, in Buckinghamshire; Godalming in Surrey; and AC on the Isle of Wight.

Aptian of Blangy (Normandy), Bize (Hautes-Pyrenees) and Haute-Savoie, in France. Possibly also from Barremian and Aptian (Bedoulian) of Provence (France).

#### Remarks

Although Parkinson was the first one to describe and figure this species (Parkinson 1822, 70-71, pl. 9: fig. 10), Mantell has been credited as the species’ author (e.g. in the Treatise by Finks et al, 2004b). It has been written that the original designation was published by Mantell (1838) in the Wonders of Geology, but the first time that Mantell figured this species is in the second volume of Wonders of Geology, 4^th^ edition, (Mantell 1839, and not 1838) without crediting Parkinson’s (1822) authorship. Nevertheless, Sherborn (1902) acknowledged this species to Parkinson (1822). Following the Article 23 of the ICZN, Principle of Priority, we credit Parkinson as the species’ author of *Barroisia anastomosans*.

Sphinctozoa basal skeleton morphology. Low growth gradient with cylindrical forms giving new ones that grow in close proximity to the previous ones. Exopores most often slightly elongated, parallel to the cylindrical forms at their fixed base. The chamber segmentation produces a zone of weakness that after the death of the individual will be often separated allowing the new osculum to have a crenulate border. Forms inhabiting shallow water.

#### Comparison

It resembles the French-Swiss species *Barroisia helvetica* (Loriol 1869), but tubular paragaster formation starts with short ridges on the wall next to the osculum. Several segment sequences, in almost cases simple, start from a common basis and rapidly increase in width; while *B. anastomosans*, on the other hand, the segment sequences form irregular, intergrown, bushy masses, and the segments have the same thickness almost everywhere. It also differs from the coeval *B. clavata* because of the lack clavate forms and transverse constrictions.

> *Barroisia clavata* (Keeping, 1883)
>
> Figures 3E-F
>
> pars 1869 *Discoelia helvetica* Loriol in Loriol and Gilliéron, 65, pl. 5: fig. 9-10.
>
> pars 1879 *Verticillites anastomans* (Mantell) in Zittel, p. 28.
>
> 1882 *Barroisia helvetica* (Loriol) in Steinmann, 165, pl. 6: fig. 5-6; pl. 9: fig.1
>
> 1883 *Verticellites clavatus* Keeping, 146, pl. 8: fig.3.
>
> pars 1884 *Tremacystia clavata* (Keeping) in Hinde, p. 176.

### Description

Sponge with cylindrical-clavate forms merged at their base (Fig. 3E). These forms are composed of superposed chambers that increase in width and have about 25-40 mm height and an approximate diameter of 8-9 mm when they emerge from the proximal base and a diameter of about 15-16 mm distally. The members of the colony may be in contact along its length. Conical paragaster. This occupies one-third of the chamber diameter (Fig. 3F). Outer surface with irregular transverse constrictions moderately developed at the chamber partition level throughout its length, but without external segmentation (Figs 3E-F). Chamber height varies, approximately from 1.5 to 3 mm. Osculum may protrude at the upper end (Fig. 3E). Endowall continuous with coarse, circular endopores in each chamber, as the previous described species. Exowall with subpolygonal exopores and triradiate on the inner layer, thickness of approximately 0.4 mm. Internal segmentations almost 0.5 mm thick.

**Fig. 5A-C.**
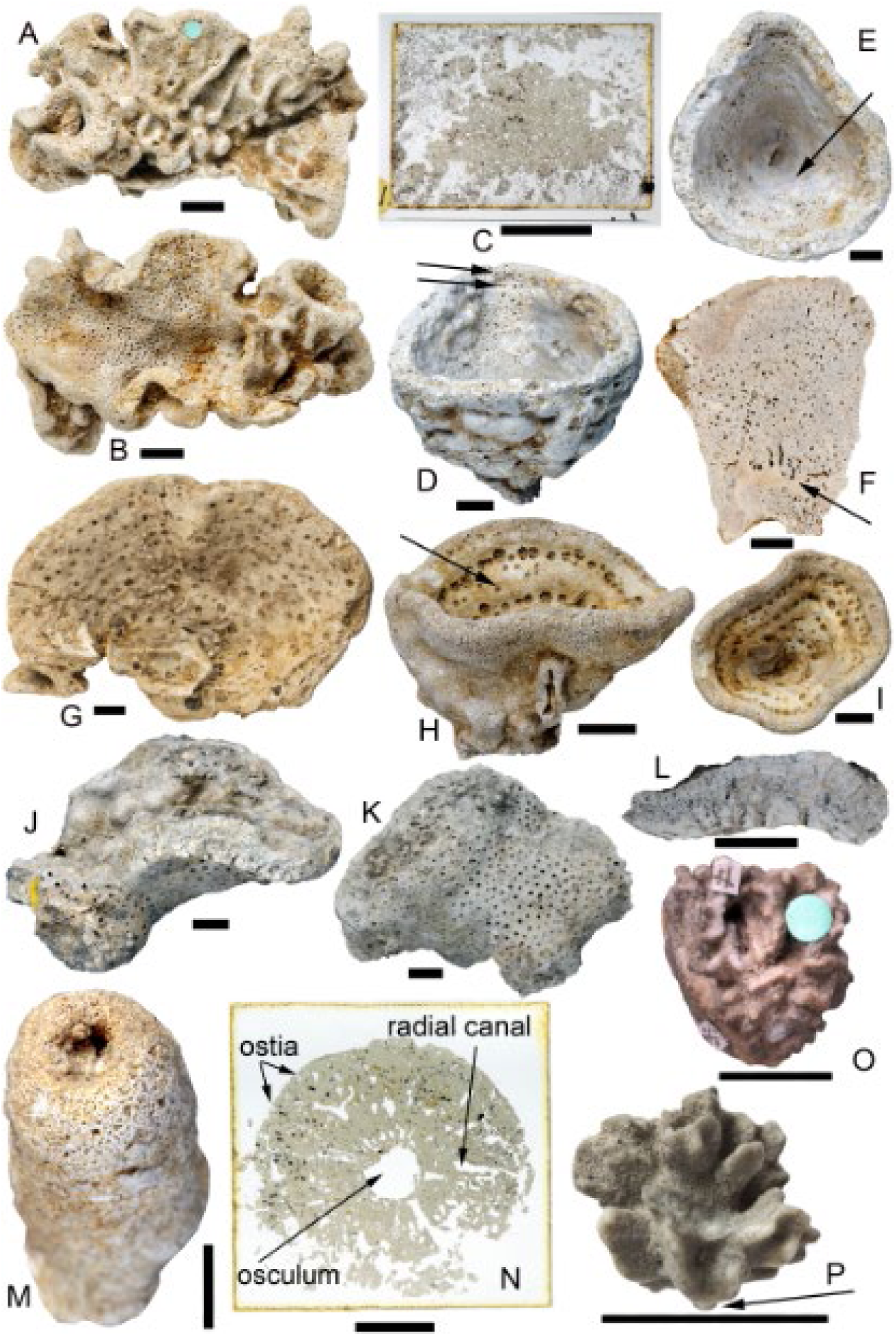
*Raphidonema porcatum* (Sharpe, 1854). A-B. Lectotype, NHMUK PI S 8578 from Faringdon, Oxfordshire. A. View of the sinuous anastomosing ridges on the external surface. B. Internal surface with oscula and pores. C. Paralectotype, NHM UK P 2181, fibres seen on a thin section. 3.3D-3F. *Raphidonema faringdonense* (Sharpe, 1854). D-E. Lectotype, NHMUK PI OR 8296. D. General view of the cup shape, with arrow on two rows of oscula, partially covered by a dermal layer. E. Internal surface of the wall with oscula, pores and dermal layer. F. Paralectotype, NHMUK S 6655, from Little Coxwell Pit, Faringdon, Oxforshire. Longitudinal section of the wall showing imperforate dermal layer (with arrow), canals and pores. G-I. *Raphidonema macropora* (Sharpe, 1854). G. NHMUK PI P 23, from Upware, Cambridgeshire, view of the concentric disposition of the apertures in the internal surface. H-I. NHM UK S 8577, from Faringdon, Oxfordshire. H. General view with external and part of the internal surfaces with annular rings alternating concentric elevations and depressions. I. View of the internal surface with the concentric rings, apertures in depressions and small pores in elevations. J-L. *Raphidonema pustulatum* Hinde, 1884. NHMUK PI P 2182, from Faringdon, Oxfordshire. J. General view with external and part of the internal surfaces with circular apertures. K. Specimen with apertures in the external surface. L. Wall section with canals seen. M-N. *Endostoma foraminosa* (Goldfuss, 1829). Lectotype NHMUK PI P 3272 and a thin section of the same specimen, NHMUK PI P 3272$1, from Faringdon, in Oxfordshire. M. General view. N. Cross section of the same specimen. O-P. *Lonsda contortuplicata* (Lonsdale, 1849), from Atherfield, Isle of Wight. O. Holotype NHMUK PI OR 46805(1), general view. P. BGS Geol.Soc.Coll.1968, general view with an arrow where the dermal layer is thicker, at the base. Scale bar 10 mm.

#### Occurrence

FSG, LGS; Aptian, Lower Cretaceous of England, from Upware, in Cambridgeshire; Faringdon, in Oxfordshire; and Little Brickhill, in Buckinghamshire.

#### Remarks

Although this species includes specimens described under *Barroisia helvetica*, it is possible to discern them from *B. clavata* for the clavate forms culminating in subglobular endings and the regular distribution of the constrictions in “*B”. helvetica*. We did not have the opportunity to study any specimen of the latter cited species microstructurally, but it seems to belong to *Tremacystia* Hinde, 1884. The constrictions of *B. clavata* seen do not follow a periodic pattern. The chambers increase in width distally, but there are zones where this is not so obvious, affecting the irregular growing of the constrictions. These data could be combined with temperature, depth and surrounding fauna to help with reconstructions of Cretaceous shallow water palaeoenvironments.

#### Comparison

It resembles *Barroisia anastomosans*, but with cylindrical-clavate forms that increase in width from base to distal end and transverse constrictions moderately developed at the chamber partition level throughout its length.

> *Barroisia irregularis* (Hinde, 1884)
>
> Figures 3G-J
>
> 1884 *Tremacystia irregularis* Hinde, 175-176, pl. 34: fig. 5.
>
> 1905 *Tremacystia irregularis* Hinde in Davey, 19.

#### Description

Colony with straight or curved cylindrical forms composed of superposed chambers of about 40 mm high and a diameter from 5 to 9 mm. Colony individuals are mostly connected (Figs 3G-H). Conical paragaster usually takes less than one-third of the chamber diameter (Fig. 3H). Outer surface mostly regularly constricted at the chamber partition level (Fig. 3G). Chamber height with an average of 1.75 mm. Osculum protrude at the upper end (Figs 3I-J). Endowall continuous with circular endopores in each chamber (Fig. 3J), almost 0.2 mm thick (Figs 3H, J). Exowall with subpolygonal exopores and triradiate on the inner layer, thickness approximately 0.4 mm. Internal segmentations approximately 0.2 mm thick.

#### Occurrence

FSG, LGS; Aptian, Lower Cretaceous of England, from Upware, in Cambridgeshire, and Faringdon, in Oxfordshire.

#### Remarks

Chambers have a regular distribution except at the flexion area in curved conical forms.

#### Comparison

It resembles *Barroisia anastomosans*, but with noticeably constricted individuals that are mostly merged and may be curved, and considerable thinner walls and chamber partitions than *B. anastomosans*. Protrusion of the osculum differs also from the previous species.

> Order STELLISPONGIIDA Finks & Rigby, 2004
>
> Family STELLISPONGIIDAE de Laubenfels, 1955
>
> Genus DEHUKIA Senowbari-Daryan, Fürsich & Rashidi, 2011

#### Diagnosis

Irregular forms with uneven folded walls which include deep grooves of different shapes. The upper part of the wall bears numerous oscula. The wall between these oscula is composed of reticulate fibre skeleton and interfibre openings.

#### Remarks

This genus includes mainly Jurassic species described from Iran, except the described species below that was previously included in *Elasmoierea* de Fromentel, 1860 (=*Elasmocoelia* Roemer, 1864, *nom. van.*).

See Senowbari-Daryan et al. (2011) for discussion.

#### Type species

*Dehukia maxima* Senowbari-Daryan, Fürsich & Rashidi, 2011.

> *Dehukia crassa* (de Fromentel, 1861)
>
> Figure 3K
>
> 1861a *Elasmojerea crassa* de Fromentel, 10, pl. 2: fig. 10.
>
> 1861b *Elasmoierea crassa* de Fromentel, 363.
>
> non 1861 *Elasmoierea crassa* de Fromentel in Loriol and Gilliéron, 70-71, pl. 5: fig. 12.
>
> 1884 *Elasmocoelia crassa* (de Fromentel) in Hinde, 176, pl. 33: fig. 11.
>
> non 1905 *Elasmocoelia crassa* (de Fromentel) sensu Hinde in Davey, 22.
>
> 2011 *Dehukia crassa* (de Fromentel) in Senowbari-Daryan, Fürsich & Rashidi, 435.

#### Description

Anastomosing forms with roughly 20 mm high and 40 mm wide, the walls of which have an approximate thickness of 4-6 mm. Circular oscula or exhalant canals arranged in rows (Fig. 3K). Oscula with an average width of 1 mm and the same distance apart each other along the crest of the walls. External fibres with trabecular microstructure, subcircular exopores, triradiate and filiform spicules.

#### Occurrence

FSG, LGS; Aptian, Lower Cretaceous of England, Faringdon, in Oxfordshire and Urgonian facies, Middle Neocomian of France, Germigney in Haute-Saône.

#### Remarks

The specimen NHMUK PI P 3273 (Fig. 3K) is the only one described (Hinde 1884: pl. 33: fig. 11) of this species from the LGS Group.

#### Comparison

Hinde (1884) included this species under the genus *Elasmocoelia* de Fromentel, 1860, but its irregular form, non-tubular, distinguishes this species of any other species of the genus *Elasmocoelia* (synonymised with *Elasmoierea*), which embraces erect laminae, mostly branching forms. On the other hand, another species included in ‘*Elasmocoelia’* and which resembles the described species is *‘E.’ tortuosa* (Loriol, 1869) that according to Davey (1905), both forms belong to the same species. But the latter has less thickness between walls and oscula are arranged in two rows abreast and not one as in *D. crassa*.

> Genus ELASMOIEREA de Fromentel, 1860

### Diagnosis

Erect, plicate, sometimes branching laminae with many vertical, exhalant canals opening mainly in single row or occasionally several abreast on upper edge. Sides of lamina may bulge around each subcircular osculum; sides of lamina covered with small, closely spaced pores. Trabecular microstructure.

#### Remarks

Some species attributed to this genus currently belong to other. Those species that remain under this taxonomic name should be revised.

#### Type species

*Elasmoierea sequana,* de Fromentel, 1860.

> *[Elasmoierea] faringdonensis* (Mantell, 1854)
>
> Figure 3L
>
> 1854 *Tragos faringdoniensis* Mantell, 229, pl. 75: fig. 5.
>
> pars 1874 *Tragos faringdoniensis* Mantell in Davey, 14.
>
> 1884 *Elasmocoelia farringdonensis* (Mantell) in Hinde, 177, pl. 34: figs 7-7a.
>
> 1905 *Elasmocoelia faringdonensis* (Mantell) sensu Hinde in Davey, 21, pl. 2: fig. 2.

### Description

Erect forms with usually fan-shaped walls. These forms are about 30 mm high and between 40 and 70 mm wide with numerous subcircular oscula without any arrangement (Fig. 3L). These have approximately 1 mm of diameter and a maximum of 1.5 mm with external borders well defined, and are connected with vertical exhalant canals. They are around 2 mm apart each other. Trabecular microstructure, with tri- and tetraradiates and slender filiform spicules. Lateral polygonal exospores.

### Occurrence

FSG, LGS; Aptian, Lower Cretaceous of England, Faringdon, in Oxfordshire.

### Remarks

*Elasmocoelia* has been synonymised with *Elasmoierea,* being the latter the senior synonym. The specific name of this taxon, *[Elasmoierea] faringdonensis,* is assigned to the genus with doubt, thus is currently in open nomenclature, which is why we have used square parentheses indicating informal identification.

This species is frequently attached to another sponge species (Fig. 3L).

### Comparison

It could be compared to *[E.] mantelli* Hinde, 1884, but this one has smaller dimensions, upper surface slightly convex, and larger osculum diameter, about 1.75-2.25 mm.

> *[Elasmoierea] mantelli* (Hinde, 1884)
>
> Figures 3M-N
>
> 1884 *Elasmocoelia mantelli* Hinde, 177, pl. 34: fig. 8.
>
> 1905 *Elasmocoelia mantelli* Hinde in Davey, 20, pl. 2: fig. 3.

## Description

Polygonal forms with expanded base, about 50 mm high and 30 mm wide. Upper surface is normally flattened, or slightly convex. With numerous circular oscula irregularly distributed (Fig. 3N). These have approximately 1.75-2.25 mm of diameter and have external borders well defined with annular membrane. The oscula communicate with cylindrical exhalant canals (arrows on Fig. 3M) which go through the sponge vertically. They do not seem to keep a pattern in their distribution on the external surface. Trabecular microstructure with tri- and tetraradiates. Fibres of about 0.15-0.30 mm thick.

## Occurrence

FSG, LGS; Aptian, Lower Cretaceous of England, Faringdon, in Oxfordshire.

### Remarks

*Elasmocoelia* has been synonymised with *Elasmoierea,* being the latter the senior synonym. The specific name of this taxon, *[Elasmoierea] mantelli,* is assigned to the genus with doubt in open nomenclature, reason why we have used square parentheses indicating informal identification.

### Comparison

It could be compared to *[E.] faringdonensis* (Mantell, 1854), but this one with larger dimensions, upper surface flattened, smaller osculum diameter, about 1-1.5 mm and fan-shaped walls.

> Genus PERONIDELLA Zittel in Hinde, 1893

### Diagnosis

Normally bush-like colony with cylindrical forms that arise from a common base, being later partly merged laterally. May be also solitary forms. The distal cylindrical end is usually rounded with central osculum connected with a deep central paragaster that runs through the form to near the base. Surface pores only regular, intertrabecular spaces. Imperforate, dermal layer present on the basal part of each branch. . Trabecular microstructure with triradites, and possibly tetradiates, including tuning-fork spicules closely intermingled. Fibres may be covered concentrically by thin layer of filiform, sinuous spicules which could line the paragaster.

### Remarks

This genus was described by Zittel (1879: p.30-31) as *Peronella*, taxonomic name already proposed by Gray in 1855 and adopted by Agassiz in 1872 for echinoderms (Hinde, 1893), therefore it was not available. Hinde (1893: p. 213) published the proposed name that Zittel offered instead of *Peronella*.

Many of the specimens belonging to this taxon have epifauna associated such as bryozoans (Fig. 3O) confirming that the parenchymal skeleton was rigid during the life of sponge.

### Type species

*Spongia pistilliformis* Lamouroux, 1821.

> *Peronidella gillieroni* (Loriol, 1869)
>
> Figure 3O
>
> 1869 D*i*scoelia *gillieroni* Loriol, 66-67, pl. 4: figs 16-17.
>
> 1879 *Peronella gillieroni* (Loriol) in Zittel, 33.
>
> 1884 *Peronella gillieroni* (Loriol) in Hinde, 169, pl. 33: fig. 10.
>
> 1905 *Peronella gillieroni* (Loriol) sensu Hinde in Davey, 17, pl. 1: fig. 3.

### Description

Bush-like colony with short cylindrical forms of about 15-30 mm high that arise from a common base reaching a colony height of 25-40 mm and a width of approximately 40-50 mm. The cylindrical forms are mostly connected (Fig. 3O). Each form has a diameter from 5 to 7 mm that ends in rounded or truncate summits. Conical paragaster with almost constant diameter that protrudes in central circular osculum of 1 mm diameter, approximately one-sixth of the diameter. Some individuals with imperforate, dermal layer at the basal part. Trabecular microstructure with triradites, and possibly tetradiates.

### Occurrence

FSG, LGS; Aptian, Lower Cretaceous of England, Faringdon, in Oxfordshire and Lower Neocomian beds of Germany, Berklingen, Lower Saxony.

### Remarks

It is characteristic that the short forms are mostly bifurcated and exceptionally trifurcous. Closely intermingled spicules normally observed in the longitudinal direction of the fiber.

**Comparison**.

It resembles *P. ramosa* (Roemer, 1839) and *P. prolifera* (Hinde, 1884). The former with smaller dimensions regarding the cylindrical forms and osculum. The latter with bifurcating forms, wider stems and oscula.

> *Peronidella prolifera* (Hinde, 1884)
>
> Figures 4A-C
>
> 1884 P*e*ronella *prolifera* Hinde, 169-170, pl. 33: figs 8-8a.
>
> 1885 ?*Peronella prolifera* Hinde in Počta, 19.

### Description

Bush-like colony (Fig. 4A) with cylindrical forms, straight or winding, on an average of 40 mm high that may bifurcate and also merge distally (Figs 4A-B) with other. These forms have a diameter of about 9-10 mm, having a central circular osculum that takes one-fifth of the total diameter (Fig. 4C), around 2 mm, connected with a deep paragaster. The distal part of the forms ends normally in rounded and occasionally inflated summit. Thick fibres from 0.2 to 0.3 mm with small triradites and tetradiates. Spicular ray length from 0.1 to 0.04 mm.

### Occurrence

FSG, LGS; Aptian, Lower Cretaceous of England, Faringdon, in Oxfordshire, and Cenomanian Korytzaner Schichten of Bohemia, Zbyslav, in Vrdy.

### Remarks

The two Bohemian specimens described by Počta (1885) coincide with Hinde’s (1884) description except the fibre thickness that is 10 times thicker. This must be a typographic mistake because the fibres cannot be thicker than the osculum.

### Comparison

It resembles *P. gillieroni* (Loriol, 1869) and *P. ramosa* (Roemer, 1839). Both with smaller dimensions regarding the cylindrical forms, osculum and fibres. The latter with spicular structure mostly obliterated and may have the lower part covered by a compact or showing irregular apertures between the fibres.

> *Peronidella ramosa* (Roemer, 1839)
>
> Figures 4D-E
>
> 1839 *Scyphia ramosa* Roemer, 11, pl. 17: fig. 27.
>
> 1844 *Scyphia ramosa* Roemer in Mantell, 256: pl. 55: fig. 5.
>
> 1861a *Discoelia ramosa* (Roemer) in de Fromentel, 9, pl. 1: fig. 5.
>
> 1861b *Discoelia ramosa* (Roemer) in de Fromentel, 362.
>
> 1884 *Peronella ramosa* (Roemer) in Hinde, 169, pl. 33: fig. 5.

### Description

Bush-like colony with short cylindrical forms of about 15-20 mm high and a width of approximately 35-40 mm. The cylindrical forms are mostly connected (Fig. 4D). Each form has a diameter from 4 to 6 mm that normally ends in rounded summits. Conical paragaster with almost constant diameter and central circular osculum of 1-1.5 mm diameter, approximately one-fourth of the diameter. Occasionally with a compact dermal layer wrapping the lower part of the stems. If this layer is not seen, showing irregular apertures between fibres. Microstructure of bounded fibres (Fig. 4E) of about 0.15 mm thick and triradites rarely seen.

### Occurrence

FSG, LGS; Aptian, Lower Cretaceous of England, Faringdon, in Oxfordshire; and Urgonian facies, Middle Neocomian of France and Switzerlnd, Germigney in Haute-Saône and Landeron, in Neuchâtel; and Lower Neocomian beds of France and Germany, Censeau, in Jura, Schöppenstedt and Schandelabe hills, in Lower Saxony.

### Remarks

It is characteristic the dense parenchymal skeleton with thin bounded fibres (Fig. 4E) of homogeneous thickness and triradites normally not seen.

### Comparison

It resembles *P. tenuis* (Hinde, 1884), differing in larger dimensions, shorter forms and more regular stem shape. This one is from Inferior Oolite.

> Genus OCULOSPONGIA de Fromentel, 1860

### Diagnosis

Sponge massive, encrusting, rounded to conical shapes with extensive, convex top. Few small, isolated circular oscula, which may be labiate, dispersed over top surface and connected with tubular canals in the interior of the skeleton, remaining surface covered with coarse pores, normally irregular, representing intertrabecular spaces. Such pores may be vertically elongate on sides. Periodic growth. Base, and in some cases sides as well, covered with an imperforate dermal layer. Microstructure trabecular, sheetlike and curve trabeculae minutely spinose at tubular interspaces, including sagittal triactines almost uniform in size; ectosome comprising sagittal and regular triactines and accompanying diactines; and sinuous filiform spicules.

### Remarks

Triactines and diactines found in Polish Jurassic species described by Hurcewicz (1975). Hinde (1893) described the sinuous filiform spicules in ectosome and triactines in central portions in a Jurassic species from South of England.

### Type species

*Oculospongia neocomiensia* de Fromentel, 1860.

> *Oculospongia dilatata* (Roemer, 1864)
>
> Figures 4F-H
>
> 1864 T*r*emospongia *dilatata* Roemer, 40, pl. 1: figs 24a-24b.
>
> 1884 *Oculospongia dilatata* (Roemer) in Hinde, 192, pl. 36: fig. 3.
>
> pars 1905 *Oculospongia dilatata* (Roemer) sensu Hinde in Davey, 23, pl. 4: fig. 1.

### Description

Low inverted cone-shape with 15-20 mm high and about 19-29 mm wide. Base covered concentrically, with an imperforate dermal layer (Fig. 4G with an arrow), can be concave or flattened. Top arched convexly with subcircular oscula irregularly dispersed over surface (Fig. 4F) and connected with tubular canals in the interior of the skeleton. Oscula usually less abundant on the superior top; approximately 1 mm diameter; slightly labiate. Exhalant canals can reach the base or only reach a short distance from the surface.

Radiating fibres (Fig. 4H) connected transversally in vertical section; thickness of 0.1-0.3 mm.

### Occurrence

FSG, LGS; Aptian, Lower Cretaceous of England, Faringdon, in Oxfordshire and “Hils-Conglomerat” (Hinde, 1884), obsolete term that in strict and regional sense it is synonymous to the Grenzlerburg Member, Salzgitter Formation, Minden Braunschweig Group, uppermost, Valanginian to lowermost Hauterivian (Erbacher et al. 2014), Lower Cretaceous of Germany, Berklingen, Lower Saxony.

### Remarks

Spicular structure in the specimens from Faringdon was not observed, but Hinde (1884) described the existence of triactines covered by sinuous filiform spicules in ectosome in the German specimens.

### Comparison

Although Davey (1905) synonymised wrongly this species with *Tremospongia pulvinaria* (Goldfuss, 1826), it differs mainly for oscula in clusters (4-6 openings per exhalant canal), these are distributed regularly, and with smaller diameter than the *O. dilatata*.

> Genus TREMOSPONGIA d’Orbigny, 1849

### Diagnosis

Sponge massive, mushroom-shaped or hemispherical, with inverted conical shape at the base that is covered by concentrically wrinkled, imperforate, dermal layer and spheroidal upper side with numerous small oscula arranged in clusters. With trabeculae and intertrabecular spaces.

### Remarks

d’Orbigny (1849) designed *Lymnorea sphaerica* as type species of *Tremospongia*, but Geinitz (1871) synonymed *L. sphaerica* with *Manon pulvinarium* (see Geinits 1871: p. 27; Senowbari-Daryan et al. 2011: p. 431). We do not agree with this synonym. Please read synonym list of *Tremospongia pulvinaria* (Goldfuss, 1826) and remarks. As Geinitz (1871) recognised both genera differ in their base, and osculum clusters may be on a higher position than the rest of the surface in *Tremospongia*.

### Type species

*Lymnorea sphaerica* Michelin, 1845.

> *Tremospongia pulvinaria* (Goldfuss, 1826)
>
> Figures 4I-J
>
> pars 1826 *Manon pulvinarium* Goldfuss, 2-3, pl. 29: figs 7a-7b.
>
> non 1845 *Lymnorea sphaerica* Michelin, 216, pl. 52: figs. 16a-16b.
>
> non 1850 *Tremospongia sphaerica* (Michelin) in d’Orbigny, 187.
>
> 1851-2 *Tragos pulvinarium* (Goldfuss) in Bronn, 61, pl. 29: fig. 1
>
> non 1860 *Tremospongia sphaerica* (Michelin) in de Fromentel, 37, pl. 2: fig. 13.
>
> 1864 *Tremospongia pulvinaria* (Goldfuss) in Roemer, 40, pl. 14: fig. 8.
>
> pars 1871 *Tremospongia pulvinaria* (Goldfuss) in Geinitz, 27.
>
> pars 1878 *Manon pulvinarium* Goldfuss in Quenstedt, 355-356, pl. 132: figs 18, 20-21.
>
> pars 1884 *Synopella pulvinaria* (Goldfuss) in Hinde, 190-191.
>
> pars 1905 *Synopella pulvinaria* (Goldfuss) sensu Hinde in Davey, 23.
>
> 2011 *Tremospongia pulvinaria* (Goldfuss) in Senowbari-Daryan, Fürsich & Rashidi, 431, pl. 4: figs. G-H; pl. 5: figs A-F; pl. 13: figs A-G.

### Description

Semihemispherical shape of about 26 mm high and 25-42 mm wide, maximum 30 and 60 mm respectively; upper surface convex with subcircular openings, from 3 to 6 units, arranged in slightly raised clusters and connected with exhalant canals (Fig. 4I). Cluster diameter of 1-2 mm and exhalant canal diameter of 0.3 mm. Openings separated from each other by 0.2 mm of fibre and separation between neighbouring clusters approximately between 5 and 10 mm depending on the specimen. Also, small irregular pores spread erratically between the fibres with 0.2 to 0.4 mm of diameter. Other parts of the skeleton with fibres between 0.15 and 0.30 mm wide. Base covered by concentrically wrinkled and imperforate dermal layer.

### Occurrence

FSG, LGS; Aptian, Lower Cretaceous of England, Faringdon, in Oxfordshire; Tourtia strata, Lower Cretaceous of Germany, Essen, in North Rhine*-*Westphalia, of Belgium, Tournay, in Hainaut; and from Upper Jurassic strata of east-central Iran.

### Remarks

Senowbari-Daryan et al. (2011) stated the clusters are not located on protuberances or nipple-like elevations as in *Mammillopora* Bronn, 1825. *T. pulvinaria* (Goldfuss, 1826) has its oscula in slightly raised clusters as Goldfuss (1826: pl. 29: figs 7a-7b) and Senowbari-Daryan et al. (2011: pl. 5: fig. A) figured hemispherical specimens with unequivocally slightly elevated clusters, differing from *Lymnorea sphaerica* Michelin, 1845 which its oscula are at the same level as the rest of the surface and is mushroom-shaped. It is obvious that this species does not belong to *Mammillopora* as there are not knoblike proper protuberances.

The Faringdon specimens are smaller than the German ones (Hinde 1884).

### Comparison

Senowbari-Daryan et al. (2011) compared this species to *T. pellisfera* Senowbari-Daryan, Fürsich & Rashidi, 2011. The latter differs in exhalant canals surrounded by thin imperforated dermal layer.

> Family ENDOSTOMATIDAE Finks, 2004
>
> Genus RAPHIDONEMA Hinde, 1884

### Diagnosis

Sponge cup or funnel-shaped with irregular or convolute outline, and relatively thin walls. Wall formed by anastomosing, tubular spaces of narrow bore, disconnected by trabeculae. Larger and straighter tubes, probably exhalant canals, pass through most of the internal wall where they run obliquely close to the wall, and open as pores with the same diameter. These tubes are arranged quincuncially. Intertrabecular spaces open as small, circular pores on both surfaces of the wall. If the inferior part of the internal surface is very thickened, it may obliterate the small pores. Microstructure with trabeculae of sinuous sheetlike or filiform forms subparallel to fibre surface made of triactines which basal part is slightly developed resembling uniaxial forms in thin sections (Fig. 4M).

### Remarks

Hinde (1884) compared this genus to *Elasmostoma* de Fromentel, 1860 and *Corynella* Zittel, 1879 (younger synonym of *Endostoma* Roemer, 1864); but it differs from them in different spicular fibres and triactines, and in their basal ray that is poorly developed.

### Type species

*Raphidonema contortum* Hinde, 1884.

> *Raphidonema contortum* Hinde, 1884
>
> Figures 4K-N
>
> pars 1874 *Manon peziza* Goldfuss sensu Davey, 10.
>
> 1884 *Raphidonema contortum* Hinde, 197-198, pl. 37: figs 2, 2a-2b.
>
> 1905 *Raphidonema contortum* Hinde in Davey, 25.

### Description

Sponge growing in convolute expansions with a general cup or funnel shape whose wall coalesce intertwining (Fig. 4K-L). Normally between 30 and 60 mm high and between 50 and 90 mm wide; wall from 4 to 8 mm thick (Fig. 4M). Internal and external surfaces, smooth in fair preserved specimens, are covered with a dermal layer of very closely arranged fibres, thinner than those of the internal part of the wall. One of these wall surfaces with circular oscula with a dimeter from 0.3 to 0.8 mm, with a random distance from each other between 0.5 and 2 mm, and associated to sinuous canals (Fig. 4M with an arrow) that normally extend at right angles into the wall. Microstructure with wall interior fibres of 0.2-0.4 mm thick, slender filiform spicules that are parallel to each other in direction of the fibre and round the limits of the canals (Fig. 4N).

### Occurrence

FSG, LGS; Aptian, Lower Cretaceous of England, Faringdon, in Oxfordshire.

### Remarks

This sponge species is frequently attached to other sponge species.

Hinde (1884) created this species with what Davey (1874: p. 10) believed to be *Manon peziza* Goldfuss, but it seems that Davey’s illustrations (1874: pl. 2 and figure on page with ‘Additional Note’ -no numbered-) refer to a different species, probably *Raphidonema pustulatum* Hinde, 1884 in case of the pl. 2. It is clear that does not correspond with *M. peziza*, but neither with *Raphidonema contortum*.

### Comparison

It is a very distinctive species, but Hinde (1884) compared to *Elasmostoma consobrinum* d’Orbigny, 1850 with which coincides in oscula diameter size and growth mode, but differs in spicular structure, thicker walls and normally larger size. It could be also compared its structural fibres to *Raphidonema porcatum* (Sharpe, 1854), but this one has characteristic sinuous anastomosing ridges on the external surface.

> *Raphidonema porcatum* (Sharpe, 1854)
>
> Figures 5A-C
>
> 1854 *Manon porcatum* Sharpe, 196, pl. 5: fig. 2.
>
> 1874 *Manon porcatum* Sharpe in Davey, 16, pl. 10.
>
> 1878 *Catagma porcatum* (Sharpe) in Sollas, 362.
>
> 1883 *Catagma porcatum* (Sharpe) in Keeping, 147.
>
> 1884 *Raphidonema porcatum* (Sharpe) in Hinde, 198, pl. 37: fig. 3.
>
> 1905 *Raphidonema porcatum* (Sharpe) in Davey, 25.

### Description

Sponge growing in convolute expansions with irregular shape or cup-shaped and sinuous anastomosing ridges on the external surface (Fig. 5A), with an average of 40 mm high, 85 mm maximum wide and 45 mm deep; wall from 4 to 5.5 mm thick. Internal surface is smooth and has numerous circular oscula of about 0.80 mm wide and irregular arrangement, apart each other between 0.2 and 1 mm; also minute pores (Fig. 5B). External surface fibres may be thinner and closer than the internal surface ones. These ones have a thickness between 0.15 and 0.3 mm (Fig. 5C). With filiform spicules.

### Occurrence

FSG, LGS; Aptian, Lower Cretaceous of England, Faringdon, in Oxfordshire, , in Cambridgeshire, and Brickhill, in Bedforshire.

### Remarks

With well-marked oscula on well preserved specimens.

### Comparison

It could be compared to *R. contortum* Hinde, 1884 regarding the spicular structure but *R. porcatum* differs in sinuous anastomosing ridges on the external surface.

> *Raphidonema farringdonense* (Sharpe, 1854)
>
> Figures 5D-F
>
> 1854 M*a*non *farringdonense* Sharpe, 196, pl. 5: figs. 5-6.
>
> 1878 *Catagma faringdonense* Sollas, 362.
>
> 1879 *Pharetrospongia farringdonensis* Zittel, 46.
>
> 1884 *Raphidonema farringdonense* (Sharpe) in Hinde, 200-201, pl. 37: figs 5, 5a-5b.
>
> pars 1905 *Raphidonema farringdonense* (Sharpe) in Davey, 25, pl. 1: upper left figure.

### Description

Sponge with a cup or funnel shape (Fig. 5D-E), with a height between 30 and 90 mm and a maximum width of 120 mm; wall from 7 to 17 mm thick. Internal surface varies with specimens. It is normally fibrous and with subcircular oscula of about 1 mm of diameter connected with exhalant canals that descend practically perpendicularly and open downward on the external surface. These oscula mostly have a random distribution, but also with a partial horizontal arrangement (Fig. 5D with arrows), apart each other between 0.2 mm and several mm; also small pores. It is partially covered by an imperforate compact dermal layer (Fig. 5E with arrow), in its most internal part and some patches close to the cup edge. External surface covered by nodose excrescences and may have thinner and closer fibres than the internal surface and also patches of the imperforate dermal layer. This dermal layer can pass through the wall (Fig. 5F). Microstructure with wall interior fibres of 0.15-0.3 mm thick (Fig. 5F), slender filiform triactines spicules that are parallel to each other in direction of the fibre.

### Occurrence

FSG, LGS; Aptian, Lower Cretaceous of England, Faringdon, in Oxfordshire, and Upware, in Cambridgeshire.

### Remarks

With well-marked oscula on well preserved specimens. Sponges found attached to others. Davey (1905) illustrated a couple of photographs of what he thought to be *R. farringdonense,* but only one of these photographs belongs to this species (Davey, 1905: pl. 1: upper left figure).

### Comparison

It could be compared to *R. contortum* Hinde, 1884 regarding the spicular structure, but *R. farringdonense* differs in lacking convolute expansions and the common existence of excrescences on the external surface. It has been reported the most similar species is the Indian Eocene *R. indica* Rigby & Mohanti, 1990, but differs mainly in its nodose excrescences and not clustered exhalant system.

> *Raphidonema macropora* (Sharpe, 1854)
>
> Figures 6G-I
>
> 1839 C*h*enendopora *fungiformis* sensu Mantell (non Lamouroux), 561: fig. 106.
>
> 1854 *Manon macropora* Sharpe, 195, pl. 5: figs. 3-4.
>
> 1874 *Manon macropora* Sharpe in Davey, 15, pl. 9: lower figure.
>
> 1878 *Catagma macroporus* (Sharpe) in Sollas, 356: fig. 1, 362.
>
> 1879 *Elasmostoma macropora* (Sharpe) in Zittel, 44.
>
> 1883 *Elasmostoma acutimargo* sensu Keeping (non Roemer), 147.
>
> 1884 *Raphidonema macropora* (Sharpe, 1854) in Hinde, 199-200, pl. 37: fig. 4.
>
> 1905 *Raphidonema macropora* sensu Hinde in Davey, 24, pl. 3: fig. 2.

### Description

Sponge funnel-shaped or cup-shaped (Fig. 5H), or forming convolute expansions, normally with a height of 50 mm (between 25 and 100 mm) and a width of 54 mm (between 23 and 110 mm); wall from 4 to 10 mm thick. Internal surface may be smooth (Fig. 5G) or with annular rings alternating concentric elevations and depressions (Fig. 5H-J), covered by a compact dermal layer perforated by circular apertures from 1.5 to 3.5 mm in diameter. These apertures have a concentric annular disposition and are situated in the depressions in case they exist. They are approximately 3-5 mm (exceptionally 0.2 mm) apart each other at the same concentric level without any pattern, and about 5 mm between two adjacent levels. They are connected with exhalant canals. Small pores are among two levels of apertures (Fig. 5H with arrow), with a random distribution and with a diameter of 0.5-1 mm. External surface is irregular and may have nodose excrescences. Microstructure with fibres from 0.13 to 0.26 mm thick forming reticulated bands. With sinuous filiforme triactines.

### Occurrence

FSG, LGS; Aptian, Lower Cretaceous of England, Faringdon, in Oxfordshire, and Upware, in Cambridgeshire.

### Remarks

The large circular apertures arranged concentrically in the internal surface are very distinctive of this species.

### Comparison

It could be compared to *R. contortum* Hinde, 1884 regarding the spicular structure, but *R. macropora* differs in more robust spicules with more random distribution, with large apertures on concentric disposition on the internal surface and possibly nodose external wall surface.

> *Raphidonema pustulatum* Hinde, 1884
>
> Figures 5J-L
>
> 1829 ?*Spongia marginata* Phillips, 168, pl. 1: fig, 5.
>
> non 1829 *Manon peziza* Goldfuss, 3, pl. 29: figs 8a-8c.
>
> 1884 *Raphidonema pustulatum* Hinde, 198-199, pl. 36: figs 8-8a.
>
> 1905 *Raphidonema pustulatum* Hinde in Davey, 24, pl.3: fig. 1.

### Description

Sponge with funnel or cup shapes (Fig. 5J), or growing with convolute expansions; normally with a height of 40 mm (between 13 and 57 mm) and width of 60 mm (between 28 and 90 mm); wall thickness from 3.5 to 13 mm (Fig. 5L). One of the wall surfaces, usually the internal one, with a compact dermal layer containing prominent circular apertures with margins normally raised, 0.5-1.20 mm diameter (Fig. 5K). These apertures connected with exhalant canals distributed regularly, apart each other equidistantly 1-3 mm. Also minute pores. The other surface is smooth with slender and closer fibres than the other surface. Fibre thickness varies from 0.1 to 0.35 mm, with filiform spicules that are parallel to each other in direction of the fibre and round the limits of the canals.

### Occurrence

FSG, LGS; Aptian, Lower Cretaceous of England, Faringdon, in Oxfordshire.

### Remarks

It is very distinctive of this species the large circular apertures with raised rim.

### Comparison

It could be compared to *R. macropora* (Sharpe, 1854) concerning the compact dermal layer with apertures, but it differs in not having concentric arrangement of the apertures and these have smaller diameter.

> Genus ENDOSTOMA Roemer, 1864

### Diagnosis

Conicocylindrical, usually simple but sometimes several basally conjoined, characterized by deep, central cloaca; principal, exhalant canals enter cloaca subhorizontally, and on top surface occur as radial grooves converging on osculum; other canals essentially intertrabecular spaces; patches of imperforate dermal layer may cover lower parts of sponge. Fibres forming bundles of mainly subparallel, extremely slender triradiates, and paratangential dermal triad tetraradiates may be present locally.

### Remarks

Mainly Hinde’s (1884: p. 181) description.

### Type species

*Scyphia foraminosa* Goldfuss, 1829.

> *Endostoma foraminosa* (Goldfuss, 1829)
>
> Figures 5M-N
>
> 1829 S*c*yphia *foraminosa* Goldfuss, 86, pl. 31: figs 4a-4b.
>
> 1844 *Scyphia foraminosa* Goldfuss in Mantell, 256: pl. 55: fig. 6.
>
> 1844 *Scyphia intermedia* Münster in Goldfuss, in Mantell, 256: pl. 55: fig. 2.
>
> 1864 *Endostoma foraminosum* (Goldfuss) in Roemer, 39, pl. 14: fig. 6.
>
> 1871 *Epitheles foraminosa* (Goldfuss) in Geinitz, 33-34, pl. 8: fig. 13.
>
> pars 1878 *Scyphia foraminosa* (Goldfuss) in Quenstedt, 351-352, pl. 132: fig. 8.
>
> pars 1883 *Corynella foraminosa* (Goldfuss) in Dunikowski, 317.
>
> 1884 *Corynella foraminosa* (Goldfuss) in Hinde, pl. 34: figs 9, 9a-9b.
>
> 1905 *Corynella foraminosa* (Goldfuss) in Davey, 16.

### Description

Conicocylindrical forms (Fig. 5M) that occur normally in solitary, but also in clusters of a few conjoined individuals. They usually have a height of approximately 45 mm and about 34 mm of diameter, being wider proximally. Deep and central cloaca of about 6 mm diameter (Fig. 5N), less than one-sixth of the diameter. There may be a differentiated layer covering the wall at the basal zone, otherwise it is possible to see ostia and canals. Endowall continuous with canal apertures present. Exhalant canals enter cloaca subhorizontally. Exowall with subpolygonal exopores. Fibres composed of minute three-rayed spicules parallel to each other in the direction of the fibre (Fig. 5N) that are combined with larger forms.

### Occurrence

FSG, LGS; Aptian, Lower Cretaceous of England, Faringdon, in Oxfordshire and Tourtia deposits, Cenomanian, of Germany, in Essen.

### Remarks

Although in cross section (Fig. 5N) seem to have filiform uniaxial spicules, they are filiform three-rayed spicules.

### Comparison

It resembles the coeval *Pachytilodia infundibuliformis* as Goldfuss (1826) illustrated it [*Scyphia infundibuliformis*]. In this case the sponge base lacks of a usual layer that *E. foraminosa* has covering ostia and canals. On the other hand, adult individuals of *P. infundibuliformis* have been described as goblet shapes with coarse, irregular pores and monoaxons besides triradiates.

> Class HEXACTINELLIDA Schmidt, 1870
>
> Order UNCERTAIN
>
> Genus LONSDA de Laubenfels, 1955

### Diagnosis

Irregular forms with numerous ridges that merge at the base; ridges and furrows alternate and may anastomose. Furrows can connect each other. Patches of compact dermal layer. Minute pores in furrows. Reticulated fibres.

### Remarks

Taxonomic name replacing *Conis* Lonsdale, 1849, due homonymy, which was created to identify a sponge from the southern basin LGS, AC of the Isle of Wight, that cannot be related to any other.

### Type species

*Conis contortuplicata* Lonsdale, 1849.

> *Lonsda contortuplicata* (Lonsdale, 1849)
>
> Figures 5O-P
>
> pars 1849 *Conis contortuplicata* Lonsdale, 63-66, pl. 4: figs 1-4.
>
> 1955 *Lonsda contortuplicata* (Lonsdale) in de Laubenfels, E86.
>
> 1961 *Lonsda contortuplicata* (Lonsdale) in Casey, 572-573.
>
> 2004 *Lonsda contortuplicata* (Lonsdale) in Finks, Reid and Rigby, 555.

### Description

Irregular (Fig. 5O) or branch-like (Fig. 5P) forms, with alternating ridges and furrows that may anastomose. Usual height between 10 and 17 mm, width between 9 and 16 mm, and approximate depth of 8 mm. Some ridges and base covered by a dermal layer that is thicker at the base (Fig. 5P with arrow); thickness varies at the same branch-like or ridge or on opposite sides. Minute inhalant pores, not seen when the dermal layer is very thick at the base. Microstructure finely reticulated with irregular meshes and bent fibres.

### Occurrence

Upper Perna Member, AC, LGS; Aptian, Lower Cretaceous of England, Atherfield and Sandown, on the Isle of Wight.

### Remarks

Irregular mode of growth that may be tuberculated. Furrows may have functioned as excurrent canals. This species has been the subject of great controversy. Casey (1961) argued this species is a hydrozoan, agreeing with the calcareous species found of shallow waters in the LGS. De Laubenfels (1955) placed this species with hyalospongea (heteractinellids) and more recently Finks et al. (2004b) with hexactinellids in order uncertain.

### Comparison

Lonsdale (1849) compared this species to the Streiberg Jurassic species *Astrospongia costata* (Münster in Goldfuss, 1829) [*Achilleum costatum*]. This one is a calcareous sponge with a hemipherical form and radiating ribs radially that meet distally differs from *L. contortuplicata* not only in form, but also in composition.

## 4. Conclusion

This comprehensive study documents 17 British calcareous sponge species and one hexactinellid from the British LGS, utilizing the latest Porifera classification and recent taxonomic research, with specimens illustrated from the Natural History Museum, London, and the British Geological Survey. These sponges are notable for their fine preservation, attributed to the loose sandy matrix and their own composition. Found abundantly in Oxfordshire, especially at the Faringdon outcrop and Little Coxwell quarries, these sponges are primarily calcareous and from littoral facies.

## Declaration of generative AI

The authors declare that no generative artificial intelligence (AI) or AI-assisted technologies were used in the creation of this article. All content, including the writing, analysis, and conclusions, was produced by the authors without the aid of generative AI tools.

## Acknowledgements

We thank Kevin Web, NHM science photographer, for the specimen macro images; and Jon Earle, NHM researcher services assistant, for helping for locating references.

